# PCBS: an R package for fast and accurate analysis of bisulfite sequencing data

**DOI:** 10.1101/2024.05.23.595620

**Authors:** Kathryn Lande, April E. Williams

**Affiliations:** The Razavi Newman Integrative Genomics and Bioinformatics Core Facility, The Salk Institute for Biological Studies, La Jolla, CA, 92037, United States

**Author notes:** Corresponding Author: Kathryn Lande - •.

## Abstract

**Motivation:** Whole-genome bisulfite sequencing is a powerful tool for analyzing chromatin methylation genome-wide, but analysis of whole-genome bisulfite data is hampered by slow, inaccurate, and inflexible pipelines.

**Results:** We developed PCBS, a computationally efficient R package for Whole Genome Bisulfite Sequencing analysis that demonstrates remarkable accuracy and flexibility compared to current tools. PCBS identifies differentially methylated loci and differentially methylated regions and offers novel functionality that allows for more targeted methylation analyses. PCBS uses minimal computational resources; a complete pipeline in mouse can run on a local RStudio instance in a matter of minutes.

**Availability and Implementation:** PCBS is an R package available under a GNU GPLv3 license at: https://github.com/katlande/PCBS and from CRAN:

https://CRAN.R-project.org/package=PCBS. Instructions for use are available at: https://katlande.github.io/PCBS/.

**Supplementary Information:** “Supplementary data are available on BioRXiv.”

## Introduction

DNA methylation plays important roles in a variety of basic biological functions such as gene splicing, transcription, and chromosomal stability, as well as in numerous disease states including cancer, autoimmune, and neurodevelopmental diseases (Jones 2012, Robertson 2005, Grolaux et al. 2022). Whole Genome Bisulfite Sequencing (WGBS) is one of the most powerful tools for assessing methylation states genome-wide. However, analyses of WGBS data are plagued by slow computational times that root from the massive sizes of whole genome data. As a result, the current analysis paradigm generally involves extracting a small number of significantly differentially methylated loci (DMLs) and/or differentially methylated regions (DMRs), and focusing exclusively on these sites for downstream analysis. This methodology risks discarding a large amount of biologically relevant information and greatly diminishes the flexibility and power granted by a whole genome dataset. Here, we introduce Principal Component BiSulfite (PCBS): a novel, user-friendly, and computationally-efficient R package for analyzing WGBS data holistically.

PCBS is built on the simple premise that if a principal component analysis (PCA) strongly delineates samples between two conditions, then the value of a methylated locus in the eigenvector of the delineating principal component (PC) will be larger if that locus is highly different between conditions. Thus, eigenvector values, which can be calculated quickly for even a very large number of sites, can be used as a score that roughly defines how much any given locus contributes to the variation between two conditions. Herein, we provide several new tools for analyzing WGBS data under this paradigm and provide a proof of concept that PCBS consistently outperforms or matches other commonly used tools in speed and accuracy metrics when used on real and simulated data .

### Results – Input

PCBS requires the sequencing depth and percent methylation for each locus in each sample, provided in a single data frame with two columns per sample. We offer a script that can convert the output of a Bismark (Krueger and Andrews 2011) alignment pipeline into PCBS input file format and also provide an example input file called *eigen* in the data of the PCBS R package.

### Results - Test Data

To test the speed and accuracy of PCBS, we used archived wildtype mouse WGBS samples from (Cole et al. 2017), representing methylation in four young and four old individuals. We used this data in combination with three simulated 24gb test genomes (1.2 million loci each) with differing amounts of true variable sites (Supp Figure 1A-B). These represent low, medium, and high variation genomes, containing 8,358, 18,774, and 37,192 “true” DMLs respectively. These DMLs are primarily assigned to “true” DMR regions of randomly assigned lengths between 100bp and 4000bp (45, 106, and 192 true DMRs in the low, medium, and high variation respectively). 1000, 3000, and 6000 stray differential loci were additionally added to each set. Differential loci were randomly assigned as either hypo- or hyper-methylated. They were also randomly assigned a 25%, 50%, or 75% modifier, representing low-, medium-, and high-intensity methylation differences between treatment and control samples.

**Figure 1:**
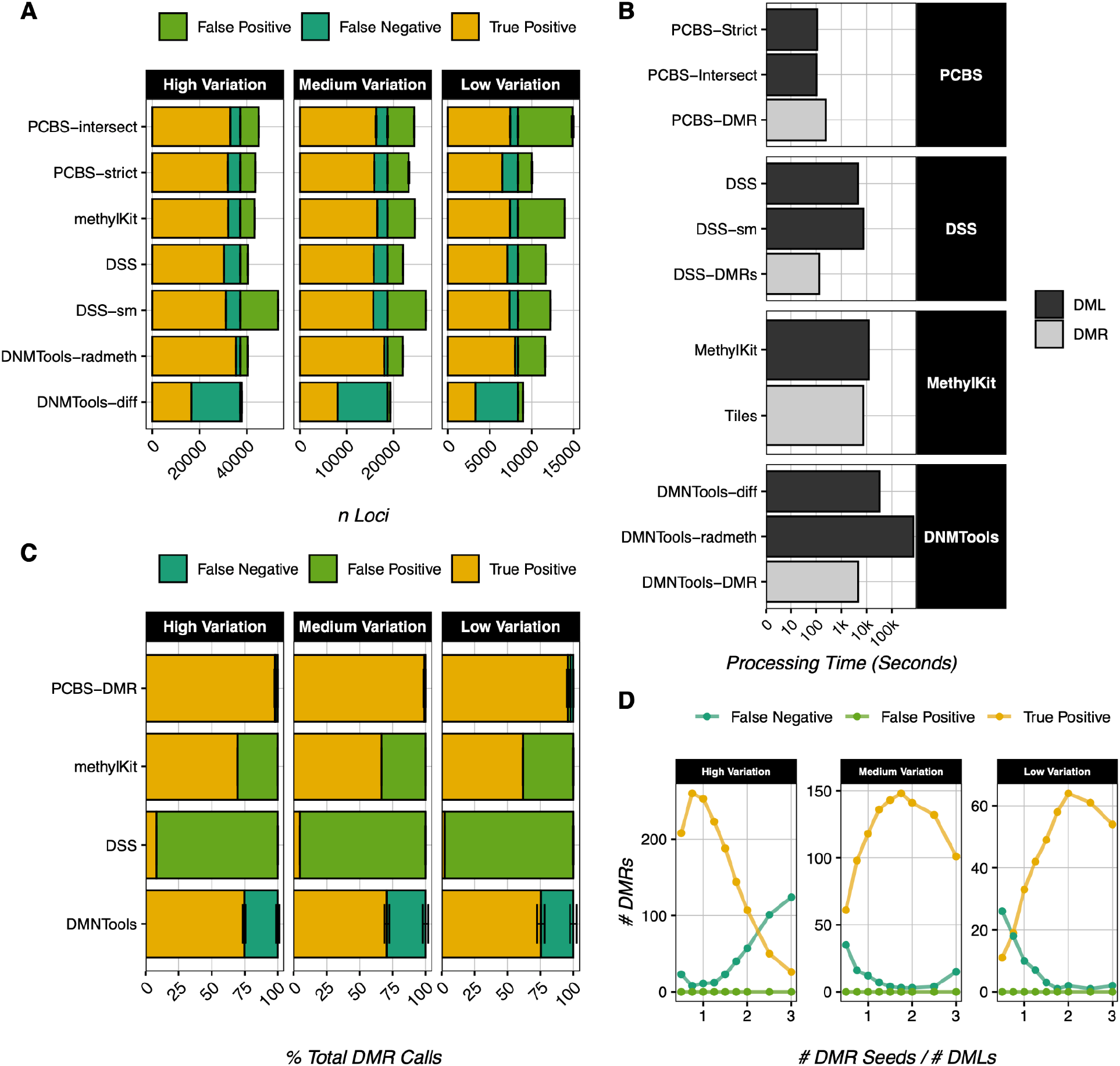
PCBS accuracy and speed testing. (a) DML calling error rates of common WGBS software and PCBS in 127 iterations of 3 treatment vs. 3 control samples for three simulated genomes of high, medium, and low variation; standard error of each call type is denoted with error bars. All DML calls use default software parameters. (b) Processing times of DML and DMR calling in common WGBS software and PCBS, on a single CPU with 8 GB RAM. All calls use default software parameters. (c) DMR calling error rates of common WGBS software and PCBS in 127 iterations of 3 treatment vs. 3 control samples for three simulated genomes of high, medium, and low variation; standard error of each call type is denoted with error bars. DMR calls are determined to be true positives if they overlap any true DMR region. All DMR calls use default software parameters. (d) PCBS DMR calling accuracy under default parameters as a function of input seed number in one iteration of 3 treatment vs. 3 control samples for three simulated genomes of high, medium, and low variation.

Per simulation, three treatment and three control samples were generated for each of the three genomes, and sites were seeded with a random percent methylation across all samples. The values of each locus’ percent methylation were pulled from a normal distribution [**μ**=0.5, sd=0.1, max=0.75, min=0.25] for NS sites, [**μ**=0.55, sd=0.1, max=0.85, min=0.35] for hyper sites, and [**μ**=0.45, sd=0.1, max=0.65, min=0.15] for hypo sites. The randomly assigned intensity modifiers were applied to each “true” site, so that there was a mix of high, medium, and low intensity percent methylation difference represented in each simulated genome.

### Results - DMLs

Most WGBS analysis software focuses on calling differentially methylated loci (DMLs) by applying statistical tests such as beta-binomial distributions (Feng and Wu 2019, Dolzhenko and Smith 2014) or logistic regressions (Akalin et al. 2012) across samples at all sequenced sites. PCBS does not do this, and opts simply to rank loci by their eigenvector score. While this does not return locus-level p-values, a simple rank cut-off performs comparably to software using the aforementioned methodology when identifying DMLs (Figure 1A), as rank order is strongly correlated to “true” DML sites in simulated datasets (Sup Figure 2A). The optimal cut-off for DML calling occurs just above the inflection point on a plot of locus rank versus absolute locus eigenvector score (Sup Figure 2B). PCBS offers two modes for estimating this cut-off. In most cases, we recommend the ‘intersect’ method, where the rank cut-off is defined as the intersection between the linear line of best fit for the highest-scoring sites (true variation), and that of the lowest-scoring sites (background noise). For very low variation datasets, we also offer the ‘strict,’ method, which functions similarly, but takes the halfway point between PCBS-intersect and the maximum rank value of the true variation line of best fit as the cut-off instead. Moreover, while DML calling accuracy is similar across software including PCBS, PCBS requires the fewest computational resources by a large margin (Figure 1B).

### Results – DMRs

DMRs are a cornerstone of WGBS analysis, and are generally defined as regions containing a number of DMLs (Peters et al. 2021, Campagna et al. 2021). Common DMR callers may look for enrichment in genomic bins (Akalin et al 2012), by merging nearby significant loci and identifying regions above a certain threshold of percent significant sites (Feng and Wu 2019), or by using hidden Markov models (Song et al. 2013). However, PCBS uses an entirely novel algorithm to identify DMRs. It works broadly by taking a user-defined rank cut-off, wherein loci above this rank are extracted as “seeds.” It then compares the median eigenvector scores of regions around these seeds against random local background regions in permutations (Supp Figure 3A). Because we expect many of the loci in a single “true” DMR region to be selected as seeds, seeds that are near each other are collapsed into single seed points at their median, and are treated as single points called “compressed seeds.” This dramatically reduces computing times. From each “compressed seed,” the algorithm expands outwards up to a maximum (user-defined) DMR size, then identifies the smallest expansion containing over 90% of the most variable sites. Following expansion, the tails of these DMRs are trimmed to remove stretches of sites with eigenvector scores similar to those of the background. Final DMR significance is calculated by comparing the rank of all DMR sites to those in a bootstrapped, randomly selected local background.

This algorithm processes relatively quickly (Figure 1B). When compared to other common DMR calling algorithms in simulated datasets, it shows the greatest accuracy by far: both in terms of total DMRs called, and at the level of individual bases within DMRs (Figure 1C, Sup Figure 3B-C). DMR calling in biological datasets also performs as expected (Supp Figure 3D).

### Results - DMR Seeds

Simulations demonstrate that this algorithm is highly resistant to false positives regardless of input seed number (Figure 1D). However, to reduce the number of false negatives, some consideration must be given when defining the number of seeds for DMR calling. If too few seeds are queried, DMRs around more weakly significant loci will be missed. Conversely, if too many seeds are included, seeds can become “overcompressed.” Overcompression occurs when seeds from multiple nearby DMRs are compressed into a single point, causing the algorithm to look for only one DMR in a region where multiple are found.

Additionally, increasing the seed number in mouse data appears to increase processing time exponentially while increasing the DMR calls logarithmically (Supp Figure 3E). Thus, for any dataset, the optimal seed number strikes a balance between including too few seeds and data overcompression. While we do offer some functionality to help safeguard against overcompression, we generally recommend using a seed number between 1-2% of the total filtered loci number to optimize the true-positive call rate against computing times.

### Results - Additional Functionality

PCBS includes additional functionality that allows users to directly query regions of interest for differential methylation by comparing the eigenvector scores of loci within a region to those in the local background. This allows users to directly assess the exact methylation levels at regions of interest in a simpler and more foolproof manner than looking for overlaps with DMRs. PCBS also offers functionality to create eigenvector score-based metagene plots of input regions.

## Conclusion

PCBS has two notable limitations. The first being that it only offers single-factor comparisons between two conditions. However, nothing about PCBS’s underlying logic precludes future development of more complex comparisons, and the speed and accuracy of PCBS in its current form is very promising.

The second limitation of PCBS is that it cannot generate significance values at the level of individual loci. However, because eigenvector rank analysis aims to examine the genome holistically, identifying differentially methylated loci is less important in a PCBS pipeline.

Moreover, a simple rank cut-off performs comparably to significance-based DML callers.

Despite these limitations, PCBS demonstrates short computational times, sensible DMR calling in archived mouse WGBS data, and unmatched DMR calling accuracy in simulated datasets. PCBS additionally introduces novel functionality that improves the flexibility of WGBS analyses. Altogether, PCBS is a powerful new option for bisulfite sequencing analysis.

## Acknowledgements/Funding Information

This work was supported by The Razavi Newman Integrative Genomics and Bioinformatics Core Facility of the Salk Institute with funding from, NIH-NIA P01: AG073084, NIH-NCI CCSG: P30 CA01495, NIH-NlA San Diego Nathan Shock Center P30: AG068635.

