## Supplemental Figures 1-3 for "PCBS: an R package for fast and accurate analysis of bisulfite sequencing data"

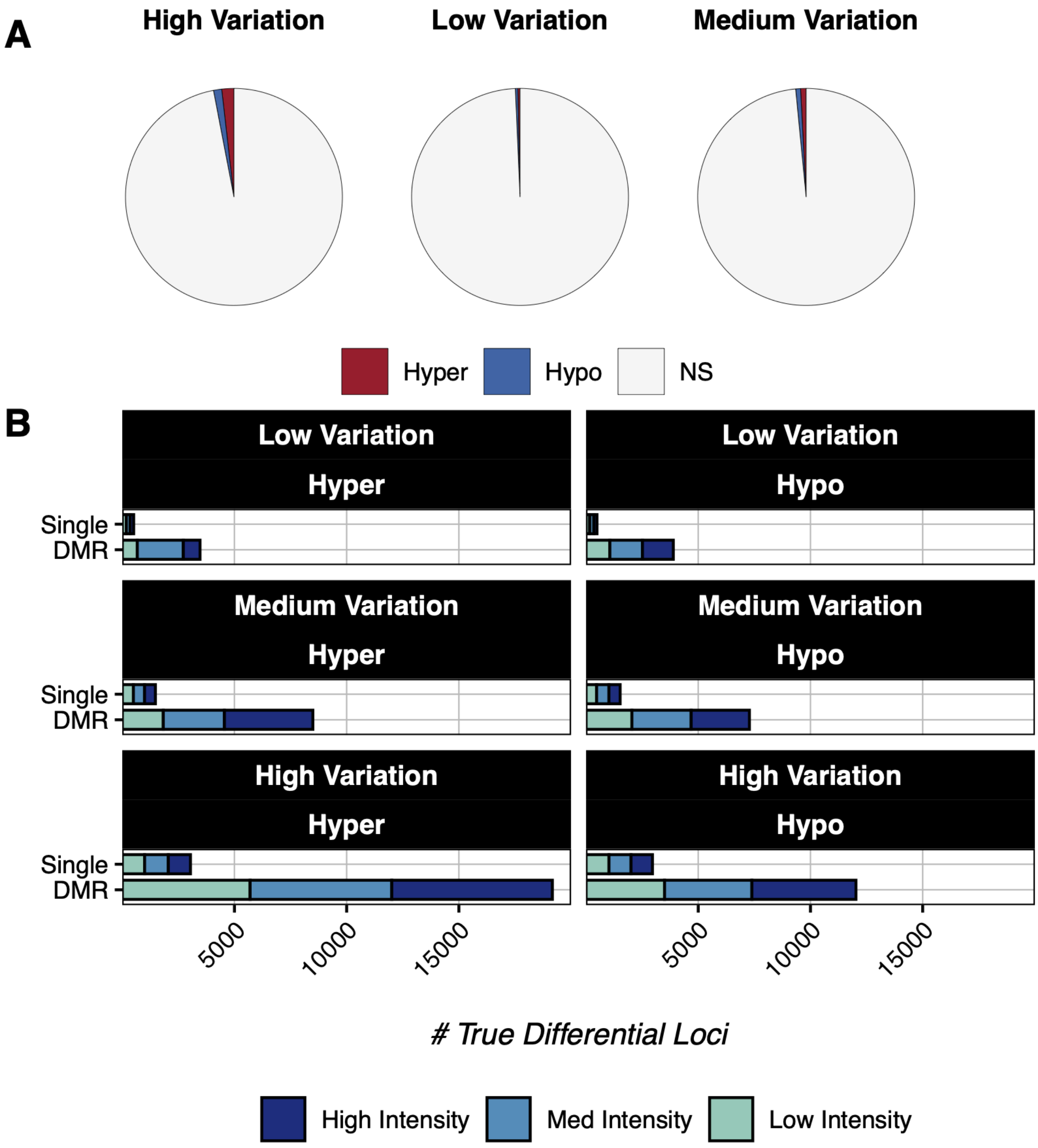


**Supplemental Figure 1: Simulated Genomes**
(a) Pie charts with the number of true hyper, hypo, and NS loci in each simulated genome of high, medium, and low variation.
(b) Barplots of the counts of high, medium, and low difference true loci in each set, divided by stray loci and loci inside defined DMR regions.

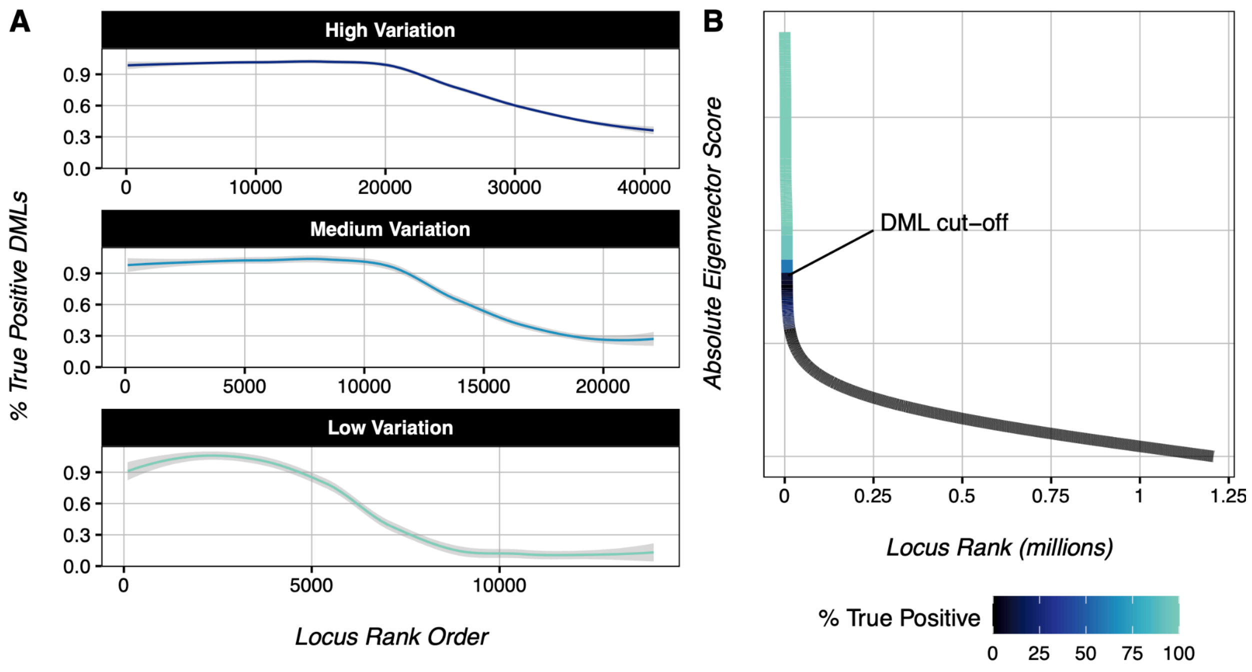


**Supplemental Figure 2: PCBS DMLs**

(a) Loess curve of % of true positive DMLs vs. eigenvector score rank order in one iteration of 3 treatment vs. 3 control samples for each simulated genome of high, medium, and low variation.

(b) Diagram of absolute eigenvector score vs. rank order, coloured by the percent of true positive loci in bins of 100 ranks. Data from one iteration of the low variation simulated genome.

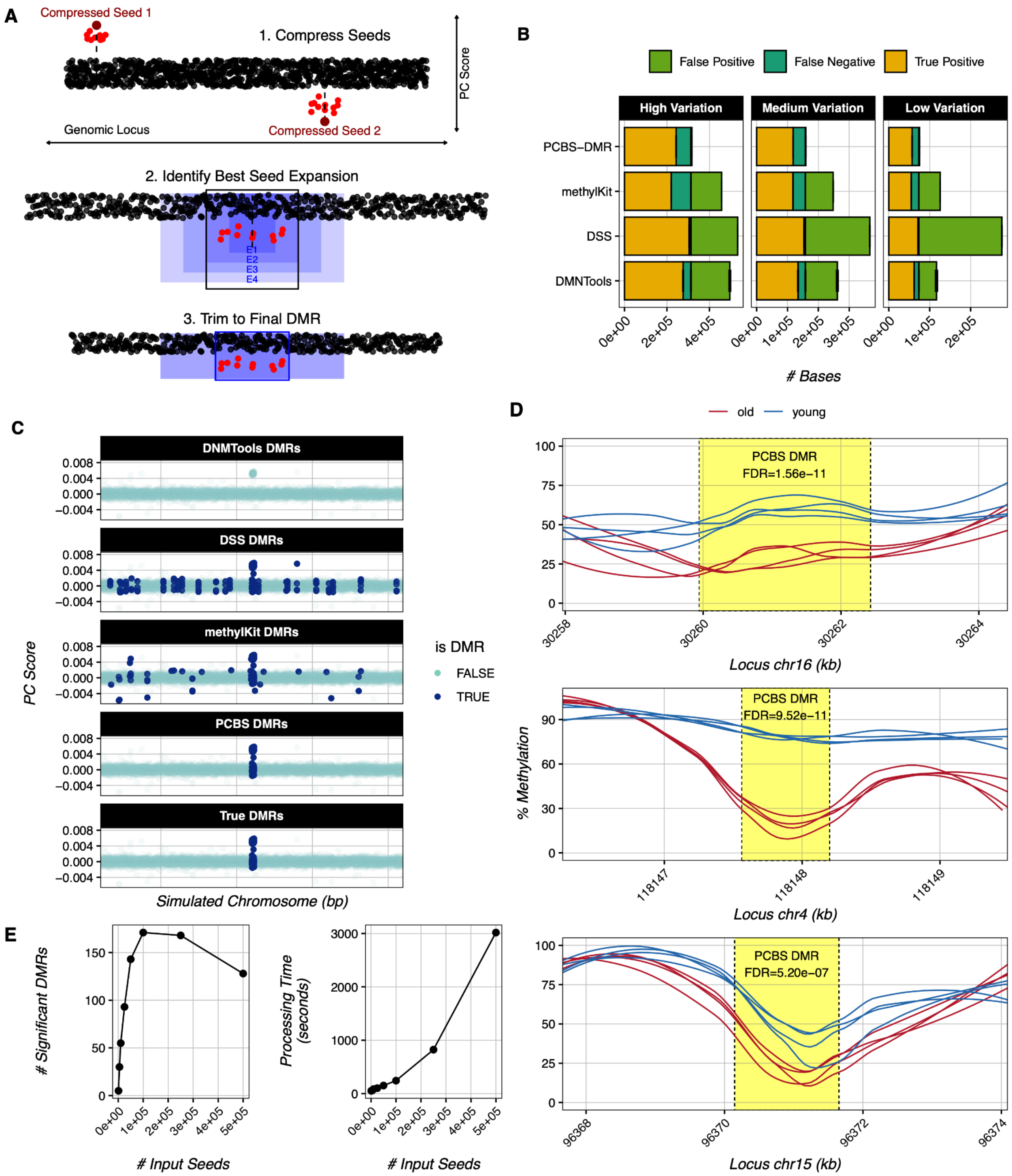


**Supplemental Figure 3: PCBS DMRs**

(a) PCBS DMR calling algorithm diagram

(b) DMR calling error rates per base pair of common WGBS softwares and PCBS in 127 iterations of 3 treatment vs. 3 control samples for three simulated genomes of high, medium, and low variation; standard error of each call type is denoted with error bars.

(c) Plots of genomic locus vs. eigenvector score of all simulated loci in a representative true DMR region, for common WGBS softwares and PCBS compared to the true DMR region.

(d) Loess curves of % methylation for four young and four old mouse wildtype WGBS samples from Cole et al 2017, overlaid with PCBS DMR calls.
(e) Plots of # significant PCBS DMR calls (FDR≤0.05) vs. input seed number, and PCBS processing time vs. input seed number.
